# Botanic Spectrum Analyser: A Deep Learning GUI for Plant Image Segmentation in Hyperspectral and RGB Phenotyping

**DOI:** 10.1101/2025.09.14.676080

**Authors:** Jason John Walsh, Levent Görgü, Emilie Jacob, Victoria Poulain, Laurent Gutierrez, Eleni Mangina, Sónia Negrão

## Abstract

Plant phenotyping systematically quantifies plant traits such as growth, morphology, physiology, or yield, assessing genetic and environmental influences on plant performance. The integration of advanced phenotyping technologies, including imaging sensors and data analytics, facilitates the non-destructive and longitudinal acquisition of high-throughput data. Nevertheless, the sheer volume of such phenotyping data introduces significant challenges for researchers, particularly related to data processing. To overcome these challenges, researchers are turning to artificial intelligence (AI), a tool that can autonomously process and learn from large amounts of data. Despite this advantage, accurate image segmentation remains a key hurdle due to the complexity of plant morphology and environmental noise. In this study, we present the Botanical Spectrum Analyser (BSA), a user-friendly graphical user interface (GUI) that integrates a modified U-Net deep neural network for plant image segmentation. Designed for accessibility, BSA enables non-technical users to apply advanced AI segmentation to RGB and hyperspectral (VNIR and SWIR) imagery. We evaluated BSA’s performance across three case studies involving wheat, barley, and Arabidopsis, demonstrating its robustness across species and imaging modalities. Our results show that BSA achieves an average accuracy of 99.7%, with F1-scores consistently exceeding 98% and strong Jaccard and recall performance across datasets. For challenging root segmentation tasks, BSA outperformed commercial algorithms, achieving a 76% F1-score compared to 24%, representing a 50% improvement. These results highlight the adaptability of the BSA framework for diverse phenotyping scenarios, bridging the gap between advanced deep learning methods and accessible plant science applications.

## 1 Introduction

Plant phenotyping is the measurement and analysis of observable plant traits, including both above- and below-ground characteristics [1]. To measure these traits, researchers have relied on traditional phenotyping techniques using destructive methodologies and manually recording data samples. However, this manual approach to plant phenotyping is time-consuming, labour intensive, and prone to human error [1]. These challenges restrict the scale and pace with which researchers can collect phenotypic data. This inherently limits progress in understanding plant growth, development, and their responses to the environment [1]. Based on these challenges, researchers have recognised the need for a more efficient and scalable approach to plant phenotyping. This has lead to the development and rapid adoption of computer vision (CV) and image-based technologies for a more non-destructive high-throughput phenotyping (HTP) approach to measuring phenotypic traits [2, 3].

HTP is a non-destructive alternative to plant phenotyping, which involves the use of advanced imaging sensors that can collect and measure plant traits without damaging the plant [4]. Recent advances in HTP technologies have revolutionised the field, providing researchers with the ability to rapidly collect measurements of plant traits in large populations [4, 5]. These new phenotyping approaches often combine science from many different areas such as robotics, CV, and artificial intelligence (AI) [4]. In particular, HTP approaches that adopt CV methodologies often use advanced imaging technologies that include RGB (red, green, blue) and spectral imaging sensors, to capture detailed plant morphology, physiology, and health information [6, 7]. Furthermore, belowground imaging systems like X-ray computed tomography (CT), magnetic resonance imaging (MRI), and rhizotron-based setups are increasingly used to non-destructively study root system architecture. Despite their value, root imaging datasets often pose additional segmentation and interpretation challenges due to soil interference, fine-scale structures, and occlusions.

RGB imaging sensors capture images in three colour bands have been widely used due to their simplicity and effectiveness in numerous phenotyping applications [3]. However, RGB sensors often lack the capability to provide detailed information on plant physiological responses. In contrast, spectral imaging sensors, which encompass visible near-infrared (VNIR) and short-wave infrared (SWIR) spectra, offer a more comprehensive view of plant characteristics by capturing hundreds of narrow spectral bands [8]. This rich spectral information enables the detection of subtle differences in plant tissues, stress responses, and other traits that are not visible in RGB images [9]. However, the complexity and high dimensionality of spectral imaging datasets poses additional challenges for image segmentation and analysis [8].

Image segmentation is the process of partitioning an image into meaningful regions, typically to distinguish objects of interest from their background. Performing an accurate segmentation of plant images is critical to extracting meaningful phenotypic information. The process is often complicated by the complex and variable nature of plant architecture, overlapping leaves, and varying lighting conditions [10, 11]. Similar challenges arise in root imaging, where fine root structures, soil interference, and low contrast can make segmentation especially difficult. However, the role of image segmentation in HTP goes beyond the separation of plant material from background [12]. Accurate segmentations are crucial for reliable trait extraction, directly impacting our understanding of plant genetics, growth, and plant stress responses [3].

Given these challenges, researchers are actively seeking reliable methods to accurately distinguish plants from their background. Traditional CV segmentation techniques, such as thresholding, are often insufficient due to the complexity of plant images. As a result, researchers have turned to more advanced segmentation algorithms, such as deep neural networks (DNNs). However, these sophisticated methods typically require technical expertise in computer science and artificial intelligence—skills that many plant scientists may not possess.

Our work addresses these challenges by presenting a novel approach to improve plant image segmentation for RGB and hyperspectral sensors. Using DNNs, we present a robust and versatile segmentation framework tailored to the complexities of plant images. The developed segmentation framework produces highly accurate segmentations of plants, which facilitates more reliable and detailed phenotypic data extraction. Furthermore, to improve accessibility and practical application in plant phenotyping, our advanced segmentation framework has been integrated into a user-friendly graphical user interface (GUI) named the Botanic Spectrum Analyser (BSA). This tool allows researchers to analyse and interpret RGB and hyperspectral plant images efficiently without the need for technical expertise. We show that BSA can be successfully used in shoot and root images, demonstrating its adaptability and performance across diverse plant imaging scenarios. Hence, accelerating phenotyping research and its applications in plant science and agriculture.

## 2 Materials and Methods

### 2.1 GUI Design and Implementation: A Comprehensive Overview

The BSA is a sophisticated Python-based application designed to facilitate the analysis of RGB and hyperspectral images [13]. This application features GUI implemented using the python library Tkinter, which is a standard GUI toolkit. The entire application is packaged into a standalone executable file, compiled and created using the python library PyInstaller. By creating a standalone executable file we ensure the ease of use across different operating system, including Windows, Linux, and MacOS systems.

#### 2.1.1 Distribution and Installation

The BSA is distributed as a single compressed ZIP file, which contains all necessary files to run the application and simplifies the installation process. The ZIP file, containing all python decencies required to run the application, bypasses the need for users to physically run an installation wizard on their local machine. To ‘install’ the application the users would simply need to extract the contents of the ZIP archive into a directory on their machine, which would create two main folders: ‘BSA-internal’ and ‘Models’.

1. BSA-internal: This folder houses all the required python dependencies that are needed to successfully run the application. These dependencies include various python packages and modules that the BSA relies upon for its operations.
2. Models: This folder contains three different deep neural network (DNN) models stored in a .h5 format. Each of these AI models is a pre-trained segmentation network, focusing on the three different species — wheat, barley and Arabidopsis — tested in our case-study analyses.

When the user extracts the full contents of the BSAs ZIP archive, several other files will be extracted to the directory. These include:

1. Labels.csv: The labels file is used for generating a categorised spectra plot of a set of hyperspectral images. It is only important for the Spectra Visualiser component of the BSA.
2. BSA logo.png & BSA logo.ico: These image files are used to enhance the user interface of the BSA by providing recognisable icons and logos on the GUI. These files are essential as their deletion or exclusion could break the BSA frontend user-interface.

Once the user has extracted all necessary files from the ZIP archive, the user can then run the included .EXE file to start the BSA application without installing a python interpreter or any additional modules. This facilitates a more self-contained nature and portable use of the application to avoid any potential challenges that the user could face through installation.

#### 2.1.2 Operational Phases

The execution of the BSA can be conceptually divided into four different operational phases: masking, configurations, analysis and visualisation. These phases are visually presented in Figure 1 and are fundamental to the BSA’s end-to-end operation. Each phase plays a crucial role in ensuring the accuracy and effectiveness of the BSAs segmentation and analysis framework. Below we discuss each phase in detail.

1. Masking: This operational phase centres around the generation of segmented images, where users can interact with the GUI to load and create their own segmentations from the list of pre-trained DNN models. The masking phase of the BSA operates as the ‘homepage’ of the application. When users start the program, the masking tools will be presented to the users. Thus, this phase also acts as the ‘starting point’ for the users analysis journey throughout the application.
2. Configurations: During this operational phase, users can configure various parameters and finetune settings to tailor their analysis in order to specific research requirements. For example, the phase focuses on selecting and uploading imaging data, providing calibration files and adjusting formulae (pre-calculated vegetation indices).
3. Analysis: In this crucial operational phase, the BSA will process all datasets uploaded by the user using the configurations set during the previous operational phase. This operation phase is straightforward and is focused purely on the BSA process and analyse and imagery.
4. Visualisation: The final operational phase of the BSA focuses on the visualisation of results generated from the analysis phase. To provide a user-friendly approach to visualising the results, BSA integrates a set of robust visualisation tools. This allows users to interpret the data through graphical representations and detailed plots.

**Figure 1:**
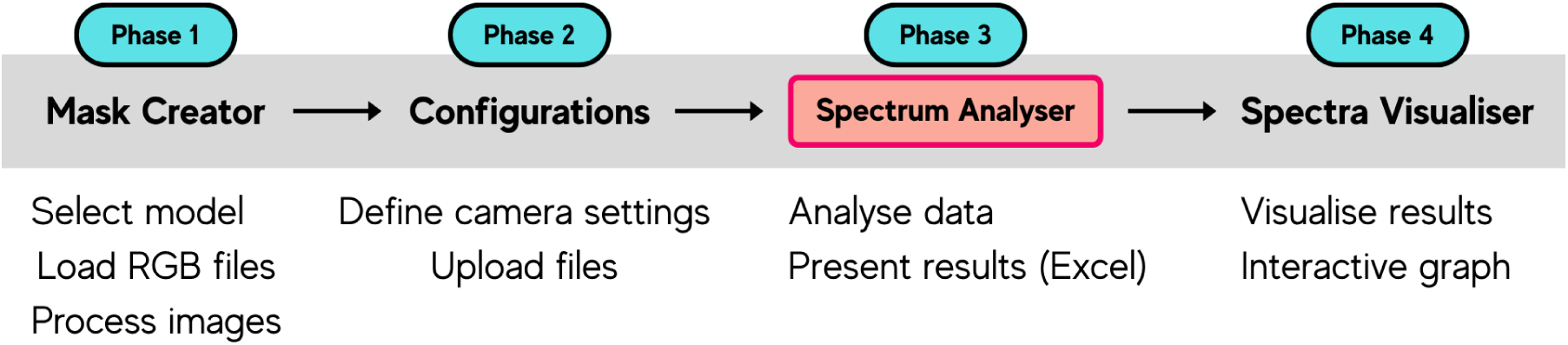
Botanic Spectrum Analyser (BSA) workflow consisting of four execution phases. Phase 1 (Mask Creator) involves selecting the model, loading RGB files, and processing images to produce a segmented image. Phase 2 (Configurations) requires defining camera settings and uploading necessary files. In Phase 3 (Spectrum Analyser), the data is analysed, and results are presented in an Excel format. Finally, Phase 4 (Spectra Visualiser) visualises the analysed results on a graph. This workflow represents the process flow for the GUI development of the BSA.

Through the seamless integration of Tkinter and PyInstaller, the BSA ensures straightforward deployment, making it accessible to users across various platforms. By structuring the workflow into these four distinct operational phases, we have prioritised user-friendliness and efficiency. This allows researchers to quickly process and analyse their data with minimal complexity. This structured approach streamlines the segmentation and analysis of both RGB and hyperspectral images, making the BSA an intuitive and powerful tool for image-based phenotyping that benefits users without advanced computational skills.

### 2.2 BSA Image Segmentation Framework

Acquiring a dataset of high-quality images is vital for plant phenotyping, as it lays the foundation for accurate analysis of morphological and physiological traits. High-resolution images enable precise image segmentation, feature extraction, and data interpretation, which ultimately enhances the reliability of phenotypic interpretation. However, poor image quality can hinder segmentation accuracy and impact downstream analysis, leading to errors in trait measurements and data inconsistencies. Image segmentation is a crucial pre-processing step in plant phenotyping, as it enables the extraction of plant pixels from the background, which ensures that only relevant features are analysed. Despite advancements in segmentation techniques, many challenges remain, particularly when dealing with complex backgrounds, variable lighting conditions, or occlusions within images each of which are common challenges inherent in plant images. Accordingly, this section focuses on the BSA’s central element—a robust deep neural network (DNN) developed for high-precision image segmentation. The BSA leverages AI-driven image segmentation to address these challenges, providing researchers with an efficient and accurate tool for analysing both RGB and hyperspectral datasets.

#### 2.2.1 Image Acquisition and Preprocessing

To appropriately train and test a DNN for image segmentation tasks, an extensive collection of high-quality imagery data is essential. Therefore, during the preliminary development phases of the BSA, we used PlantScreen™ System from Photon Systems Instruments (PSI, Drásov, spol. s r.o., Czech Republic) to gather a robust dataset for training and testing the BSA’s image segmentation framework. The complete information of the system and image capture is detailed below in the case study analysis section. In summary, the PlantScreen™ System included specialized RGB, VNIR, and SWIR imaging sensors, each housed in a dedicated chamber. This enclosure ensures that the plants are imaged under consistent and controlled lighting conditions and are free from any external interference. To ensure that each enclosure is connected, a conveyer belt runs through each chamber to transport each of the plants for imaging. Given the imaging suites enclosed nature, we capitalised on this to minimise potential disruptions by ensuring that each plant was exposed to identical environmental conditions during all imaging procedures. Shoot images were captured from the ExHIBiT barley collection which comprises 363 spring barley accessions [14]. After imaging collection, we focused on randomly selecting 100 images from each imaging sensor dataset (RGB, VNIR and SWIR) to establish the ground truth for image segmentation tasks — an essential reference for evaluating segmentation accuracy. Each ground truth images was manually segmented using the open-source image editing software GIMP to ensure a reliable and precise segmentation, an example of which can be seen in Figure 2. These manually segmented images served as the baseline for evaluating the performance of the BSA’s segmentation framework. Furthermore, for the hyperspectral images only, we extracted a single RGB image slice for each image with dimensions of 500 × 500 px. Through a series of experiments with 3-dimensional imagery, we came to the conclusion that by instead working with RGB slices we reduce the computational complexity and minimise the resource demands.

**Figure 2:**
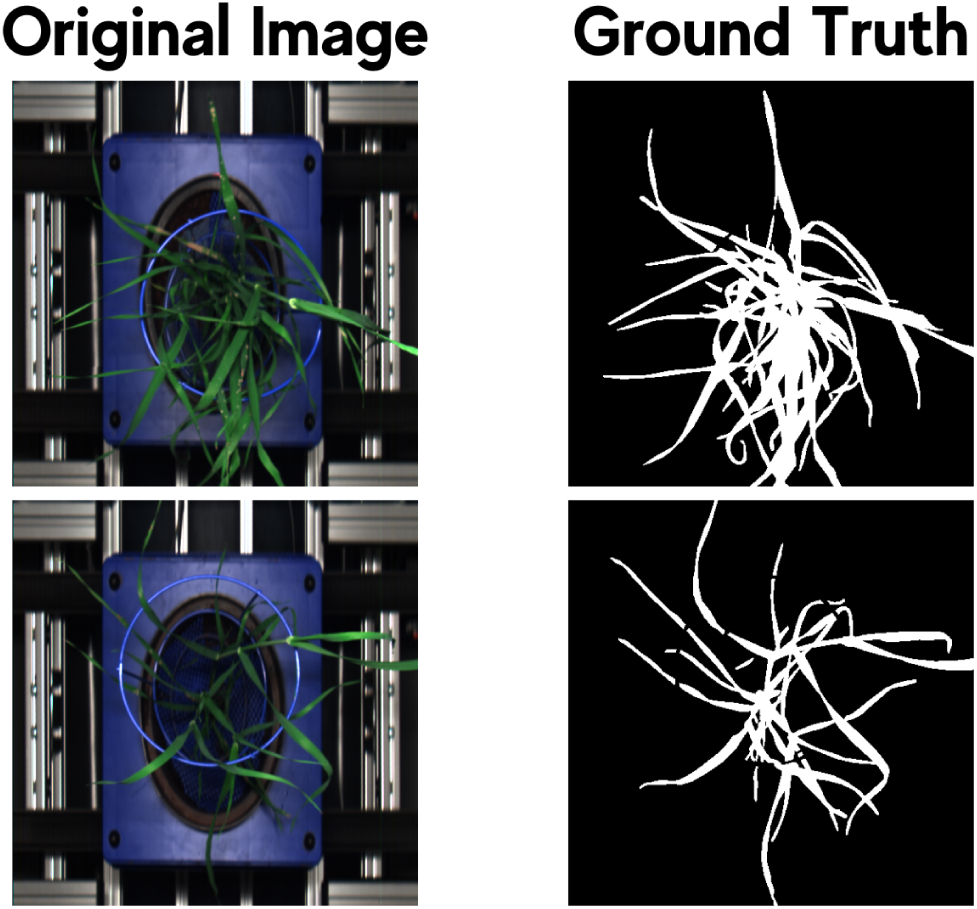
Example of original plant images (left) and their corresponding ground truth segmentation masks (right). The original images show top-down views of plants with varying leaf structures, while the ground truth masks highlight the segmented plant regions in white against a black background. These annotations serve as reference data for evaluating segmentation performance in plant phenotyping applications.

Given the relatively small size of our dataset we could not efficiently train any DNN without a substantial amount of data. Thus, to overcome this challenge, we applied aggressive data augmentation to our dataset in order to enhance the diversity and robustness of each of the collected images. Data augmentation involves applying a range of image transformations such as shifts, rotations, flips, and other geometric alterations to create a suite of synthetic images. Although the primary goal of this process was to artificially expand the dataset, we also implemented it to simulate variations found in real-world scenarios, thereby enhancing the model’s ability to generalize across different conditions and plant structures. For each of the datasets used, the data augmentation process was carefully applied to each of the randomly chosen images, resulting in a substantial increase in dataset size. After data augmentation had been applied, each dataset comprised approximately 1,000 images for training. An example of the data augmentation pipeline used can be observed in Figure 3.

**Figure 3:**
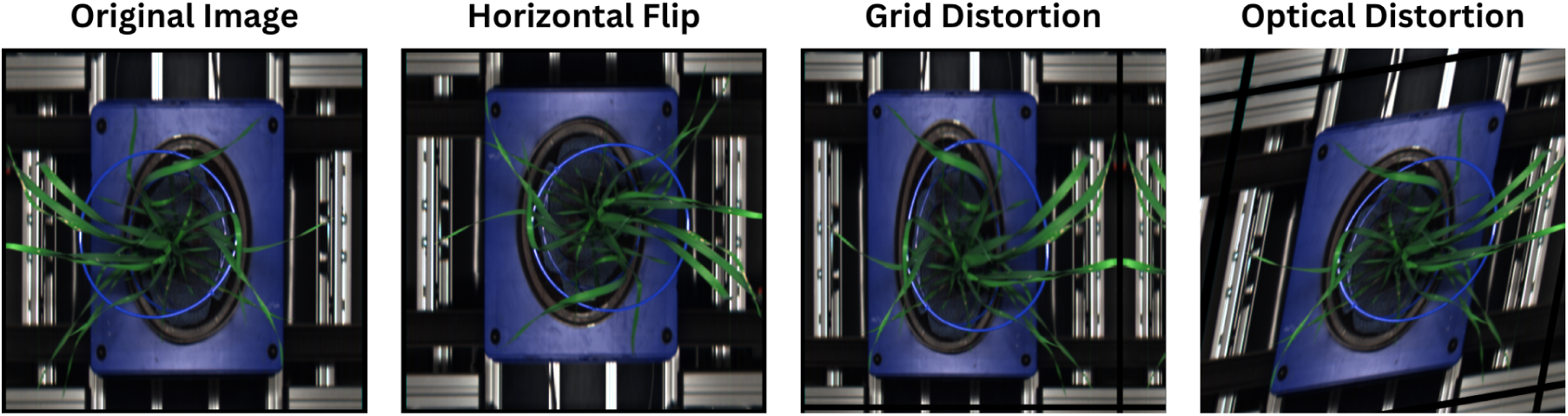
Illustration of data augmentation techniques applied to plant images. The original image (left) is modified using different transformations: horizontal flip, grid distortion, and optical distortion. These augmentations enhance model robustness by introducing variations in the dataset, aiding in generalization for plant phenotyping tasks.

**Figure 4:**
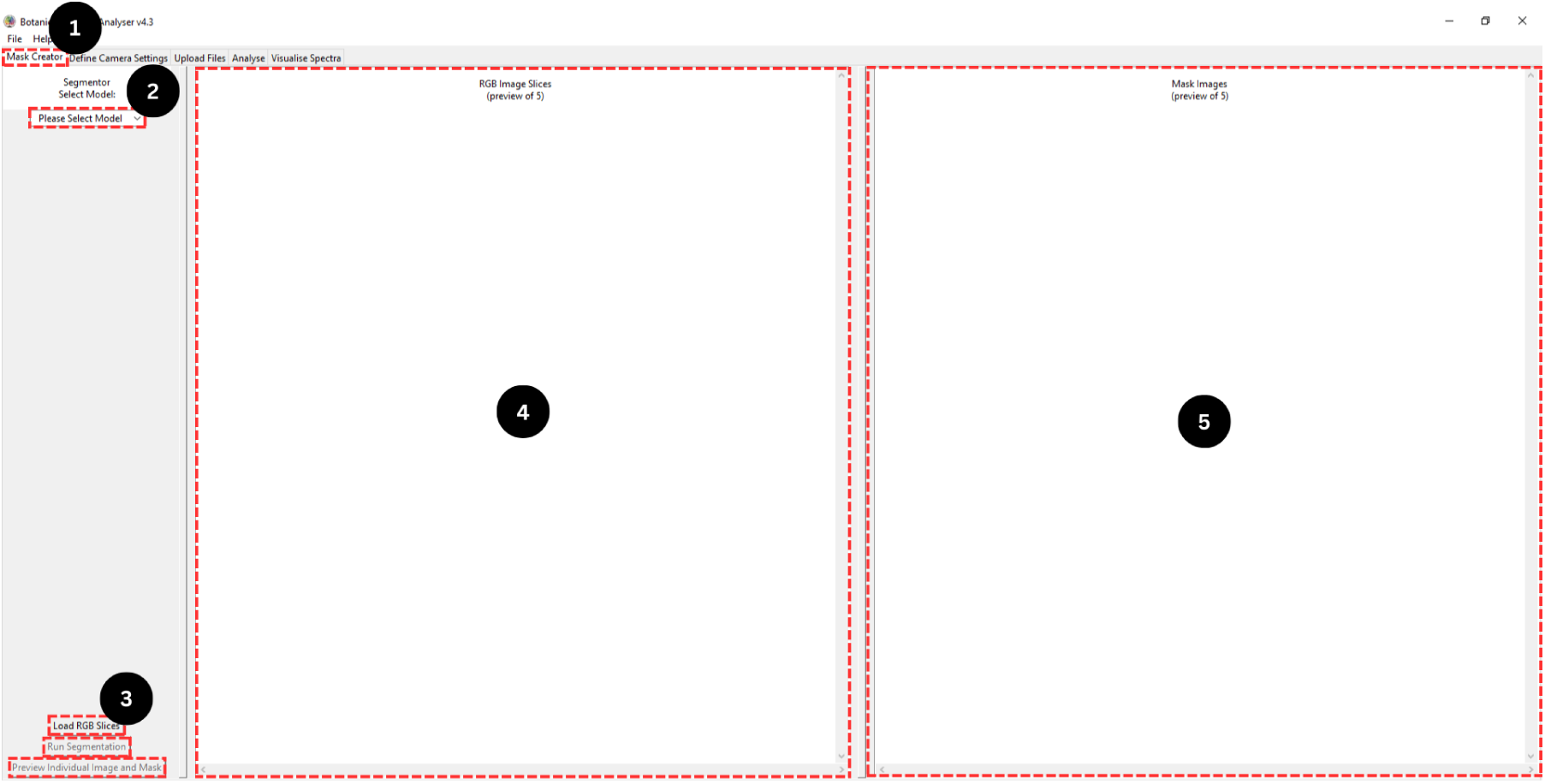
Graphical user interface (GUI) of the Botanic Spectrum Analyser (BSA) with key features labelled. (1) The ‘Mask Creator’ tab, used for generating binary segmentation masks from hyper-spectral images. (2) The model selection drop-down for choosing a pre-trained or user-imported segmentation model. (3) Buttons for loading RGB slices, running segmentation, and previewing results. (4) The left panel displaying a preview of loaded RGB slices. (5) The right panel, which appears after segmentation, showing a preview of generated masks.

**Figure 5:**
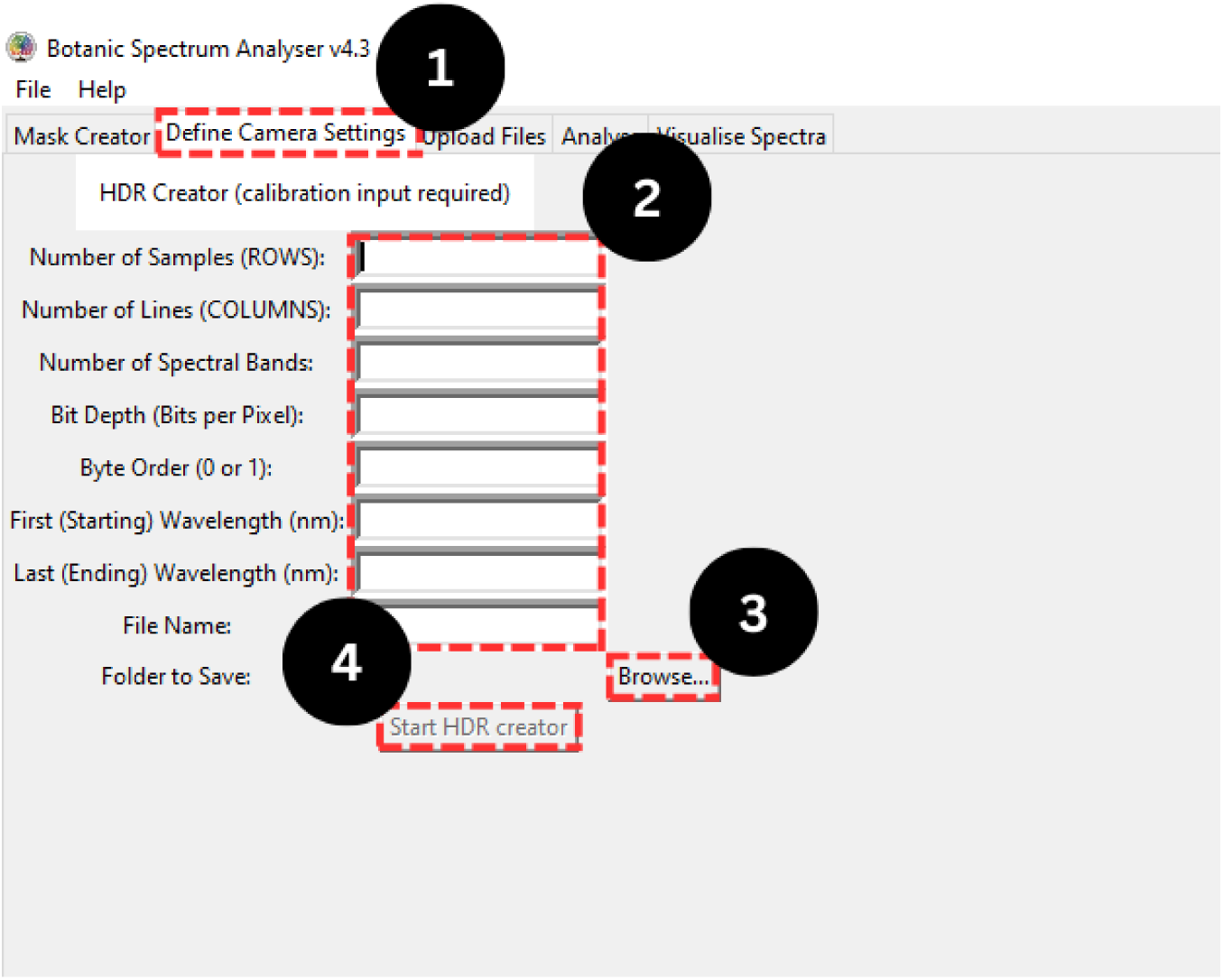
The ‘Define Camera Settings’ tab in the Botanic Spectrum Analyser (BSA) GUI, used for creating high dynamic range (HDR) files. (1) The tab location for accessing the HDR Creator tool. (2) Input fields for specifying HDR parameters, including spectral bands and wavelength range. (3) The ‘Browse’ button for selecting a save location. (4) The ‘Start HDR Creator’ button to generate an ENVI-format HDR file after completing the required inputs.

**Figure 6:**
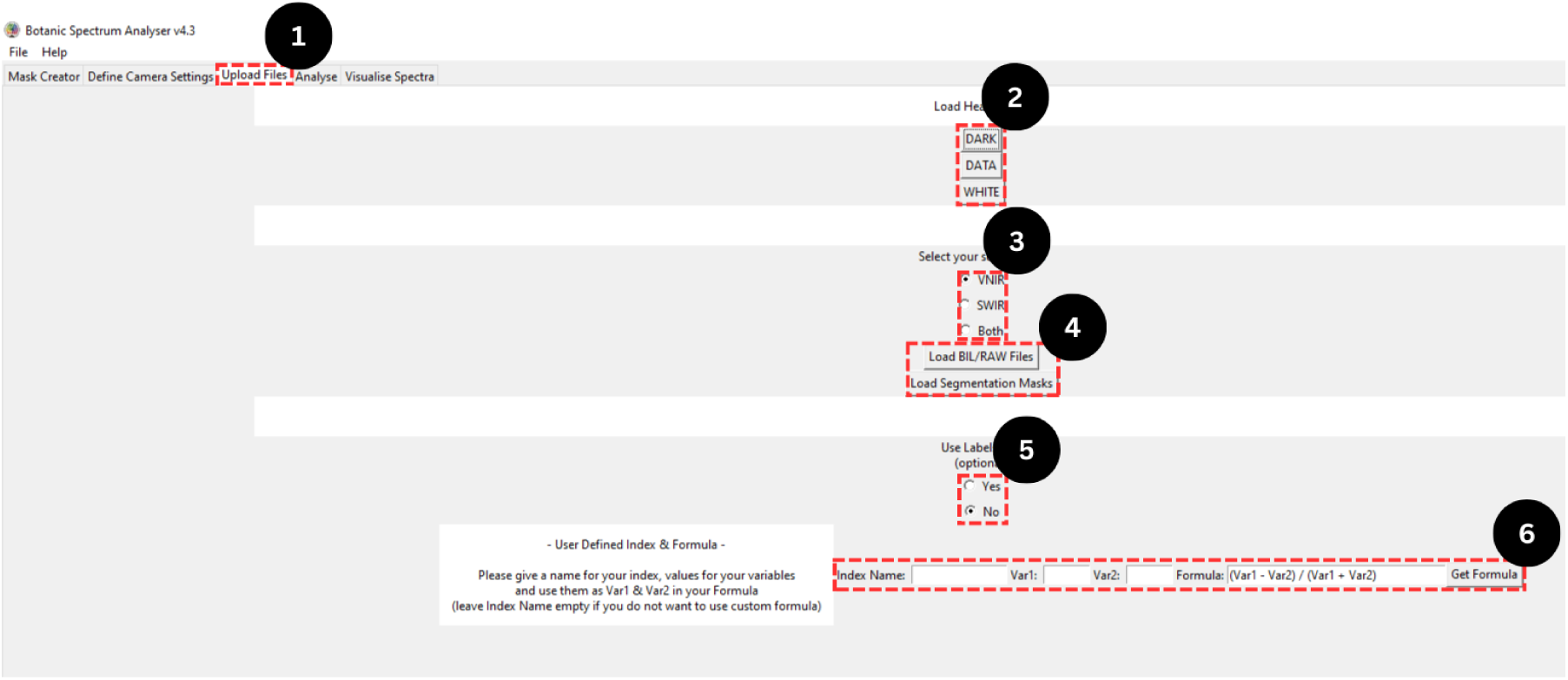
The ‘Analyser’ tool in the Botanic Spectrum Analyser (BSA) GUI. (1) The ‘Upload Files’ tab, which serves as the entry point for data analysis. (2) Buttons for uploading HDR files, including data and calibration files. (3) Sensor selection options for specifying the imaging type. (4) Buttons for uploading hyperspectral image datasets and segmentation masks. (5) An optional label selection feature for classifying images into groups. (6) A custom metric injector for defining user-specific vegetation indices, supplementing the pre-calculated indices.

#### 2.2.2 Segmentation Network Overview and Training

To produce accurate segmentations for each of the imaging datasets, we selected the popular DNN architecture U-Net based on its track record of efficiency and accuracy in image segmentation tasks, similar to that of this study. To adapt and improve the baseline U-Net architecture to our plant images, we modified the network to reduce both its computational load and increase its efficiency, resulting in faster training and increased accurate predictions. Our modifications optimised the network, in order to process a large amount of images more quickly, enabling a real-time analysis of each imaging dataset (RGB, VNIR and SWIR). Despite an extensive list of modifications made to the networks architecture, the network still follows an encoder-decoder structure. This architectural design helps retain important spatial details for accurate identification of plant features from each image. The encoder block of the network, extracts key patterns from the input images, while the decoder reconstructs them at their original scale (500 x 500 px) to enhance segmentation accuracy. To improve segmentation performance, we incorporated batch normalization and dropout layers, which reduce the overall risk of overfitting and make the model more adaptable to new images. The final layer of the network assigns each pixel to a specific class (0 or 1), enabling precise segmentation of plant images.

To evaluate the network after training, the BSA segmentation framework was evaluated using precision, recall, intersection over union (IoU), and the Dice coefficient — metrics widely employed in previous studies to assess segmentation accuracy in plant phenotyping and biomedical imaging (Zhang et al., 2019; Li & Wang, 2018). The predictions would then be generated on a previously unseen test dataset to ensure an unbiased assessment of the generalisability of the model.

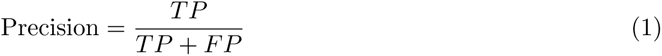

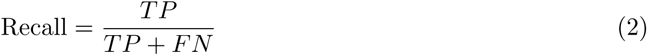

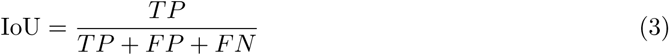

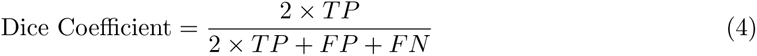

where:

- *TP* = True Positives (correctly predicted positive cases)
- *FP* = False Positives (incorrectly predicted positive cases)
- *FN* = False Negatives (positive cases that were incorrectly predicted as negative)

Following testing, the segmentation quality of each prediction was quantified using a suite of performance metrics which included the mean pixel accuracy, F1-score, Jaccard index, and recall, which were computed for segmentation image.

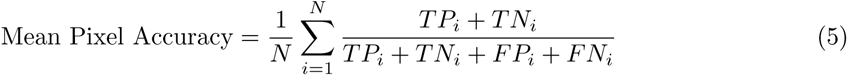

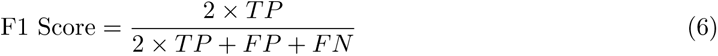

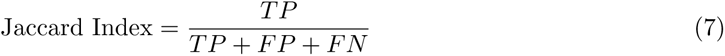

where:

- *N* = Total number of classes (or instances, depending on your context)
- *TN_i_* = True Negatives for class *i* (correctly predicted negative cases)

Furthermore, while the current model was trained on datasets generated in controlled environments, future iterations of the BSA could integrate transfer learning or pretraining on publicly available plant image datasets to further enhance generalizability across diverse species and field conditions.

### 2.3 BSA Functionalities and Workflow

The BSA offers a wide range of functionalities designed to support various research needs. The design and development of the GUI followed a structured workflow to ensure an intuitive user experience. This approach aimed to optimize user interaction and minimize potential challenges that may arise. In this section, we outline the purpose, features, and integration of the GUI within the broader system. To effectively illustrate these functionalities, we provide examples in each subsection, which demonstrate how users can interact with the BSA to achieve their research objectives.

#### 2.3.1 User Interface and Navigation

The BSA includes a user-friendly GUI with a tab-based design, ensuring intuitive and easy navigation. Throughout the user’s interaction with the BSA, text-based directions and warnings are provided to guide users on the required inputs or actions. Additionally, the interface uses colour changes (red to green) and button state changes (usable to disabled/blurred) to indicate progress through the program’s four execution phases.

The BSA GUI consists of several tabs, each of which aligns with one of the four execution phases.

1. Mask Creator (Initial Execution Phase): This is the first tab of the BSA and serves as the starting point of the application. Here, users generate binary segmentation masks from their provided images. The tab features a two-window design, with a side panel containing essential buttons for interacting with the mask creator. The central display dynamically updates to show both the original image and its segmented version, providing a preview of the generated results.
2. Define Camera Settings (Second Phase): The second tab of the BSA provides users with text boxes and buttons to create high dynamic range (HDR) files. This tab is optional and not required for dataset analysis but serves as an additional functionality for users. Since HDR ENVI (Environment for Visualizing Images) files are a mandatory requirement for BSA, this tab was designed specifically to generate HDR files in the ENVI format. Users who do not already have ENVI HDR files are encouraged to use this feature to ensure compatibility with the system.
3. Upload Files (Second Phase): The third tab of the BSA plays a key role in the configuration phase of the application by providing users with a comprehensive set of options to upload essential files, including images, calibration files, HDR files, and segmentation binary masks. Together with the second tab, it forms the configuration phase. This step is mandatory and must be completed before proceeding with spectral analysis.
4. Analyse (Third Phase): This fourth tab contains little functionality but provides the necessary buttons to allow the users to start the analysis process. After feeding the required inputs and configurations into the application from the ‘Upload Files’ tab, users are granted access to run the analyser, which will process the data and present the results in a tabular format similar to an Excel spreadsheet.
5. Spectra Visualiser (Final Phase): The fifth and final tab of the BSA provides the necessary tools for visualising the analysis results through graphical representations. This tab displays a scatter plot showing the mean spectra of each provided hyperspectral image embedded within the GUI. Users can save the generated plot in their preferred format using the available buttons. Since the spectral plot is based on the spectral bands of hyperspectral images, this tab is not compatible with RGB images and requires hyperspectral input.

Each of these tabs forms the workflow of the BSA, designed with user experience in mind. Interactions with the GUI should follow a logical sequence, ensuring a smooth and efficient workflow for data processing and analysis.

#### 2.3.2 File Handling and Data Requirements

The BSA relies on efficient file loading and structured data management to maintain optimal performance and prevent system overload. Given the large size of hyperspectral imaging datasets and to facilitate a streamlined workflow, users must specify a directory for data loading, which enables an organized approach to handling imaging datasets. Once the directory has been chosen, the BSA will not load all files simultaneously as this would exceed memory capacity and degrade the systems performance. To mitigate this challenge, we purposely designed a sequential data-loading approach for the BSA, which requires that files are processed incrementally rather than in bulk. This method optimises memory usage, prevents system slowdowns, and maintains efficient data handling regardless of the dataset size that the user uploads.

Beyond efficient file handling, proper data input is essential for ensuring accurate analysis. To use the BSA for hyperspectral image analysis, users must provide a set of required files before the process can begin. Specifically, the BSA requires three types of hyperspectral image files: a data file (hyperspectral image), a white calibration file, and a dark calibration file. Additionally, each hyperspectral file must be accompanied by its corresponding HDR file, which contains critical metadata such as spectral and spatial dimensions.

Along with the imagery data and HDR metadata, users must also supply binary segmentation masks for the dataset they are working with. These files are mandatory for accessing the analysis portion of the BSA. However, if a user intends only to use the mask creator, they need to provide RGB images in PNG or JPG format, without the requirement for hyperspectral data.

#### 2.3.3 Mask Creator

The mask creator is a key tool within the BSA, designed to enhance the segmentation accuracy of plant images captured using RGB or hyperspectral imaging sensors. A binary segmentation mask is an image that uses two colours, typically black and white, to distinguish different regions of interest (ROI) within an image. In plant science, these masks are essential for isolating plant material from the background, enabling more precise analysis of plant health, growth, and other characteristics.

The mask creator was developed with several key objectives in mind:

1. User-friendly design: The BSA mask creator was built to provide an intuitive and accessible tool for generating high-quality segmentations of plant images, whether they are in RGB or hyperspectral formats.
2. Deep learning integration: To support non-technical users, the tool enables the application of deep learning networks for image segmentation without requiring specialized expertise or retraining, making advanced image analysis more accessible to the wider plant community.

Since the mask creator can segment both RGB and hyperspectral images, it supports BIL (Band Interleaved by Line) and RAW image formats, making it compatible with a wide range of hyperspectral imaging sensors. However, the mask creator does not work with 3D hyperspectral cubes. Instead, the mask creator works with 2D RGB slices of each hyperspectral image, which optimizes both the speed and efficiency of the segmentation framework. If the user wishes to segment hyperspectral images using the mask creator, they must first extract RGB slices from their hyperspectral BIL or RAW files. These slices, derived from the visible spectrum segments of the hyperspectral cube, are essential as they contain the necessary colour information for accurate mask creation.

With their RGB slices extracted and uploaded into the BSA, the ensuing step for users involves loading a suitable segmentation network to use for mask creation. Given the users’ specific research task, the user should select a suitable DNN model from the drop-down menu provided in the mask creator tab. Although the BSA provides pre-trained segmentation architectures, the mask creator also provides the option of importing a custom DNN model, which allows users to tailor the segmentation process to their specific needs.

Upon selecting the appropriate DNN model, users can initiate the image segmentation process by simply clicking on the start button. This action triggers the application of the loaded DNN, where segmentation tasks are executed in the back-end of the BSA. The chosen DNN will process the users loaded RGB slices and identify and extract all the plant material from each image.

The final outcome of the mask creator is the generation of a highly accurate binary segmentation mask of each uploaded image that are automatically exported to a specified local directory on a user’s computer in PNG format. This feature not only simplifies the workflow but also saves valuable time, allowing researchers and analysts to concentrate on in-depth data exploration and analysis. This automated segmentation framework significantly reduces the time and effort required for manual segmentation, providing a non-technical solution for accurate and efficient image segmentation.

#### 2.3.4 HDR Creator

The HDR creator in the BSA is an optional tool designed to assist users in generating HDR files in ENVI format for their hyperspectral images. Also known as header files, HDR files are essential in hyperspectral image analysis as they contain critical metadata required for accurate spectral interpretation and processing. Without HDR metadata, several challenges can arise when handling hyperspectral images:

1. Loss of essential spectral information: Metadata such as the number of bands, spectral wavelengths, and spatial resolution is crucial for accurately interpreting hyperspectral data. Without this information, the data may be misrepresented, leading to incorrect analyses and conclusions.
2. Incompatibility with image-processing algorithms: Many hyperspectral analysis algorithms rely on metadata to function correctly. Missing HDR information can prevent these algorithms from processing the data properly, leading to errors, misalignment, or suboptimal results.

Thus, the HDR creator provides users with a simple and efficient way to generate their own HDR files. As an optional tool, it is not part of the BSA’s core workflow but serves as a supplementary feature for users who need to create HDR files in ENVI format.

To generate an HDR file, users must input the required metadata — such as the spectral and spatial dimensions of their hyperspectral image — into the provided text boxes, specify a save location, and initiate the creation process. The output is a new text-based HDR ENVI file, which can then be used during the analysis phase of the BSA.

#### 2.3.5 Hypercube Analysis

Beyond the mask creator, the BSA’s analysis phase stands as another key component, offering a robust platform to seamlessly analyse hyperspectral images. To analyse these images, we designed BSA to automatically calculate a series of vegetation indices that are crucial for assessing plant health and physiological status. The BSA achieves this by utilising the required files that users must upload during the configuration phase. By providing the HDR files, the application automatically extracts the necessary wavelength information, allowing the full spectral range of the user’s hyperspectral files to be accommodated regardless of the spectral image being used (e.g., VNIR and SWIR).

To provide the user with greater control over the analysis process, we added two optional features to the BSA. The first optional feature provides users with an opportunity to include a labels file. A labels is simply a CSV file which categorises images (hyperspectral data) into specific classes, e.g., stress vs. non-stress; disease vs. control. Although the labels file is not a mandatory requirement, the addition of this file provides the option to create a targeted spectra visualisation of the images and reveal the mean spectra per class.

The second optional feature we implemented is the addition of a custom metric injector, which allows users to compute their own custom vegetation indices. By specifying the band number from the desired wavelength for calculation, users can create and calculate personalised indices tailored to their specific research requirements. Currently, the BSA calculates seven different vegetation indices, six of which are designed for VNIR spectral images (images ranging from 300 - 900 nm) and one for SWIR images (images ranging from 900 - 1700 nm). These vegetation indices and their formulas that the BSA uses for calculation are provided in Table 1.

**Table 1:** All available vegetation indices pre-calculated via the Botanic Spectrum Analsyer (BSA). In total, eleven vegetation indices were provided through the analyser of BSA. Each of these was chosen based on their popularity and widespread used in plant phenotyping with spectral imaging sensors. The formulas provided for each vegetation index are the exact formula used by BSA analyser to calculate each vegetation index based on the type of imaging sensor used (visible near infrared or short wave infrared).

| Vegetation Index | Sensor | Formula | Reference |
| --- | --- | --- | --- |
| Normalised Difference Vegetation Index | VNIR | $(R_{800} - R_{670}) / (R_{800} + R_{670})$ | [15] |
| Photochemical Reflectance Index | VNIR | $(R_{570} - R_{531}) / (R_{531} + R_{570})$ | [16] |
| Plant Senescence Reflectance Index | VNIR | $(R_{680} - R_{501}) / R_{750}$ | [17] |
| Structure Insensitive Pigment Index | VNIR | $(R_{800} - R_{445}) / (R_{800} + R_{680})$ | [18] |
| Normalized Difference Red Edge Index | VNIR | $(R_{790} - R_{720}) / (R_{790} + R_{720})$ | [19] |
| Water Potential Index | VNIR | $(R_{665} - R_{715}) / R_{715}$ | [20] |
| Normalized Difference Water Index | SWIR | $(R_{860} - R_{1240}) / (R_{860} + R_{1240})$ | [21] |
| Moisture Stress Index (MSI) | SWIR | $(R_{1600} / R_{820})$ | [22] |
| Normalised Burn Ratio (NBR) | SWIR | $(R_{860} - R_{2210}) / (R_{860} + R_{2210})$ | [23] |
| Cellulose Absorption Index (CAI) | SWIR | $(R_{2000} - R_{2100}) - (R_{2100} + R_{2200})$ | [24] |

#### 2.3.6 Spectra Plotting and Visualisation

Upon completing the analysis phase in the BSA, a spectra plot is automatically generated, which provides a visual representation of the analysis results for each hyperspectral image. A spectra plot displays the mean spectra of each hyperspectral image uploaded to the BSA. The visualisation represents the average spectral response across all analysed pixels within each hyperspectral image. To create the spectra plot, the BSA calculates the mean spectra for each individual hyperspectral image provided. The resulting data is then visualised in the spectra visualizer tab using the python package matplotlib. By default, this visualisation is rather simple and plain, displaying only the mean spectra for each hyperspectral image. However, the visualisation can be significantly enhanced with the optional use of a labels file. When a labels file is provided, the BSA performs additional calculations, incorporating class-based distinctions for each hyperspectral image. For example, if the labels file categorises images into two classes, such as ‘stress’ and ‘non-stress,’ the BSA will compute three distinct spectra plots: one representing the overall mean spectra of each image, a second for the ‘stress’ category, and a third for the ‘non-stress’ category. If the labels file contains more than two classes, the BSA will generate separate mean spectra plots for each identified class.

## 3 Case Study Analysis

To rigorously evaluate the performance of the BSA’s segmentation and analysis framework, we selected four distinct datasets, each containing RGB or hyperspectral imaging data acquired using VNIR and SWIR imaging sensors across three different plant species — barley, wheat and Arabidopsis. These datasets were chosen to assess the framework’s adaptability to varying spectral ranges, species characteristics, organ specificities, and imaging conditions.

Each dataset presented unique challenges, such as variations in spectral reflectance, structural differences among species and organs, and potential noise from environmental factors. By testing across diverse datasets, we aimed to ensure the robustness and generalisability of the BSA in accurately segmenting plant material and extracting reliable spectral information for phenotypic analysis.

### 3.1 Case Study 1: Hyperspectral VNIR Sensors for Nitrogen Estimation in Wheat

Liu et al. (2020) investigated the application of hyperspectral imaging sensors for the estimation of nitrogen levels in wheat plants. In the study, the authors used both hyperspectral cameras and non-imaging spectrometers to measure nitrogen content in wheat leaves non-destructively. Through a series of experiments, the authors evaluated different sensor setups and explored the use of VNIR and SWIR spectra to assess their accuracy in nitrogen quantification. Their research provides valuable insights into the potential of hyperspectral sensing technologies for HTP nitrogen monitoring in wheat.

#### 3.1.1 Datasets Used

In the study, Liu et al. (2020) used data collected from five of the most widely grown wheat varieties: Gladius, Axe, Scepter, Corack, and Yitpi [25]. These wheat varieties exhibited different morphological and spectral features at the growing stage of 40 to 60 days. The plants were grown in a controlled greenhouse environment and subjected to four different nitrogen treatments (25, 50, 100, and 200 mg N/kg), with ten biological replicates per treatment. The resulting dataset consisted of hyperspectral measurements captured at three time points over the course of the study, totalling 543 valid samples after outliers and errors were removed.

#### 3.1.2 Experimental Design and Implementation

In their work, Liu et al. (2020) employed both imaging and non-imaging hyperspectral sensors across VNIR and SWIR spectra [25]. The hyperspectral data were collected using three distinct setups: a non-imaging spectrometer for point-based measurements and two hyperspectral cameras—one for whole-plant imaging and another for individual leaf scanning. The spectrometer operated across a wide spectral range (350 to 2500 nm), while the VNIR and SWIR cameras captured more targeted spectral bands. All data were collected in a controlled environment using automated phenotyping platforms to ensure consistent lighting and measurement conditions. To improve data quality, pre-processing techniques such as Savitzky–Golay filtering and spectral band resampling were applied before performing partial least squares regression (PLSR) to estimate nitrogen content in the plants [26, 27].

### 3.2 Case Study 2: Hyperspectral SWIR Sensors for Waterlogging Stress Response in Barley

We conducted a study at University of Picardy Jules Verne (UPJV), using hyperspectral SWIR imaging sensors, to investigate the waterlogging stress response in barley. The dataset was derived from the ExHIBiT collection, which includes a wide range of European barley varieties [28]. A selection of these barley samples was imaged using a state-of-the-art system (PlantScreen™ from Photon Systems Instruments – PSI). This imaging system captured detailed SWIR hyperspectral data, providing insights into the physiological responses of barley under waterlogged conditions. This approach allowed for non-invasive monitoring of plant stress, specifically targeting the spectral range associated with water content and stress indicators in plant tissues.

#### 3.2.1 Datasets Used

A representative core collection of 230 genotypically and phenotypically diverse accessions coming from the ExHIBiT population [28] was grown in the automated PlantScreen™ Modular Facility from October 2021 to February 2023. In each of the six different HTP runs that were performed, up to 176 plants were cultivated in controlled environment (day/night photoperiod of 16H/8H and temperatures of 19°C/15°C). The experimental design consisted in growing side-by-side two plants from the same accession, each one being submitted to different watering modalities: waterlogged or non-stress control conditions (split-plot design). To evaluate the diversity of waterlogging response and ability to recover among this core collection [28], the waterlogging stress was applied at 3rd leave-stage during 14 days, before a recovery phase of 7 days [29]. Raw SWIR-images and corresponding binary masks were randomly used among these 21-days’ time-series to train the plant mask definition model of the BSA, regardless of the barley accession (landrace, old-or elite-cultivar) or the watering modality applied (control or waterlogged). The heritage accession Golden Promise and the established commercial malting cultivar RGT Planet were included as reference lines in these experiments as well as in the panel of training-images.

#### 3.2.2 Experimental Design and Implementation

Daily images acquisition was performed to dynamically monitor the morphometric and high-resolution spectral signature of the aerial surface of barley plants, with RGB, VNIR and SWIR sensors. The projected shoot area (PSA) was calculated as the sum of segmented pixels in RGB images and compared to the shoot dry mass weighed at the final day of imaging, to assess the accuracy of the image-based phenotyping. Hyperspectral imaging was carried out in the range of visible and near-infrared spectrum (VNIR, 380 – 870 nm) and shortwave infrared spectrum (SWIR, 930 – 1670 nm), with two different cameras, displaying a spectral resolution of 0.8 nm and 2 nm, respectively. Indices correlated with physiological and biochemical characteristics, such as water content, were calculated by the Plant DataAnalyzer software (version 3.3.16.3, PSI), providing an image analysis of plants on a pixel by pixel basis which was averaged for each plant. The resulted dataset was a great source to improve segmentation using thresholding method for instance. The limitations previously faced were mainly caused by water reflection in waterlogged plant. They were significantly overcome by AI-driven segmentation to obtain a robust plant mask-definition.

### 3.3 Case Study 3: RGB Imaging Sensors for Trait Extraction in Plant Phenotyping

Minervini et al. (2016) presented a finely annotated dataset and proposed image segmentation techniques to advance automated image-based plant phenotyping [30]. The authors introduced a dataset with images of Arabidopsis and tobacco plants, providing pixel-level annotations for plant structures and image segmentation tasks. The study aimed to improve the accuracy of phenotyping tasks such as leaf segmentation, plant detection, and biomass estimation through advanced computer vision techniques. The research by Minervini et al. (2016) provides essential resources and image-based phenotyping methodologies for HTP applications in controlled environments.

#### 3.3.1 Datasets Used

The study by Minervini et al. (2016) used datasets containing time-lapse images of two model plants: *Arabidopsis thaliana* and *Nicotiana tabacum* (tobacco) [30]. Plants were grown under controlled conditions, with variations in lighting, moisture, and plant age to introduce diverse segmentation challenges. The dataset includes over 40,000 images, captured at several developmental stages, providing both raw images and binary masks. For Arabidopsis, images were collected over 7 weeks using different camera setups, capturing the plants in various conditions. For tobacco, images were taken over 30 days using a robotic imaging system designed to simulate diverse lighting and growth conditions. This annotated dataset allows for detailed analysis of plant structure, making it suitable for training and evaluating ML models in leaf segmentation, plant detection, and morphological feature extraction.

#### 3.3.2 Experimental Design and Implementation

In their study, Minervini et al. (2016) focused on image-based segmentation of plants to facilitate trait extraction tasks relevant to plant phenotyping [30]. The dataset includes binary and instance-based segmentation masks, with binary masks representing plant vs. background and instance-based masks differentiating individual leaves within each plant. Each mask was binarized, allowing semantic segmentation for plant area analysis. The researchers used ML algorithms, such as active contour models and support vector regression (SVR), to segment leaves and estimate plant biomass. These segmentation techniques were tested against various environmental conditions (e.g., overlapping leaves, shadowed areas) to ensure robustness.

To further improve their image segmentation accuracy, Minervini et al. (2016) applied preprocessing techniques including colour normalization and Savitzky–Golay filtering [30]. For model evaluation, metrics such as Dice Similarity Coefficient and Mean Intersection over Union (mIoU) were also used. These approaches enabled accurate segmentation of plants and leaves, supporting downstream phenotyping tasks such as biomass estimation, leaf counting, and trait extraction. This work notably highlights the potential of computer vision in advancing HTP applications in image-based plant phenotyping but also offers an invaluable dataset for training and benchmarking image segmentation models.

### 3.4 Case Study 4: Hyperspectral SWIR Sensors for Trait Extraction in Root Phenotyping

Within the scope of the research project conducted between UPJV and UCD (previously mentioned in the case study 2), a new experimental approach was set up in January 2025 to explore the use of SWIR spectra recorded from roots. This part of the study aimed to correlate root-derived hyperspectral signatures with nutrient content obtained via elemental analysis in plants subjected to waterlogging stress.

#### 3.4.1 Datasets Used

A rhizobox-adapted waterlogging protocol was optimised at UPJV to study the root responses of barley lines from the ExHIBiT core-collection and plants grown in a controlled environment (day/night photoperiod of 16H/8H and temperatures of 19°C/15°C). A 5 days-waterlogging stress followed by a 7 days-recovery was imposed at 3 leaves-stage plants in this pilot experiment, according to the described methods by [29]. Moreover, the design of custom-made rhizoboxes dedicated to sampling roots at different key time points were obtained by including a nylon membrane with a mesh size of 30 µm, confining the roots in a 2D plane, preventing them from growing in the soil compartment of the rhizobox, while allowing water and solutes to flow to the roots. This method enabled gathering roots non-soil contaminated, making them suitable for hyperspectral imaging phenotyping before destructive sampling for elements’ analyses or microscopy. To evaluate the segmentation performance of the GUI-based segmentation, ten SWIR images acquired on roots of the reference cultivar RGT Planet cultivated under both control and waterlogging conditions were analysed.

#### 3.4.2 Experimental Design and Implementation

To identify new imaging markers related to root traits and their plasticity under waterlogging conditions, a suite of available tools was employed at UPJV. Hyperspectral imaging was performed with the previously characterized automated PlantScreen™ Modular Facility, a phenotyping platform designed for analysing the shoots and thus being most likely inefficient in terms of image segmentation at the root level. Excavated roots from rhizoboxes were gently removed from the nylon membrane and placed on a discriminant black background to be imaged using the SWIR camera (930 – 1670 nm; section 3.2). The specific purpose was not to assess the morphological characteristics of the root system but rather to explore the reflectance features extraction from roots of waterlogged and control plants. The objectives were to compare these two watering conditions and investigate the ability of barley plants to modify their root element content after being exposed to waterlogging-induced hypoxic conditions and subsequent recovery. The generated dataset clearly demonstrated the effectiveness of the BSA’s masking algorithms on root images, showcasing its potential to provide an accurate mean spectrum for below-ground phenotyping.

## 4 Results and Discussion

### 4.1 BSA Segmentation Framework Delivers High Level Segmentation Results

To train and evaluate the BSA segmentation framework, we conducted a series of experiments using our PSI imaging suite dataset. Each imaging dataset was used to develop a tailored DNN optimised for its unique characteristics at segmenting RGB and hyperspectral images. Consistent with best practices in DL-based image segmentation research [31, 32], we adopted an 80:20 data split, ensuring robust training while preserving a portion of the dataset for independent validation.

By evaluating the BSA’s initial segmentation performance, we observed that the framework demonstrated significant improvements over both traditional and DL-based segmentation algorithms. We benchmarked against PlantCV [33], a widely used open-source plant imaging analysis tool that applies vegetation indices such as the popular Excess Green (ExG) formula to enhance plant features for segmentation [34]. Our method consistently outperformed the python library in terms of segmentation accuracy. This result aligns with findings from literature, which showed that DL algorithms consistently surpass traditional rule-based segmentation algorithms in capturing the finer details in plant images [35, 36].

Figure 7 visually illustrates these performance differences, presenting the original hyperspectral image, the ground truth, our BSA segmentation framework’s (U-Net model) prediction, and the overlay of the segmentation mask. The strong alignment between the predicted and ground truth masks underscores the robustness of our proposed approach.

**Figure 7:**
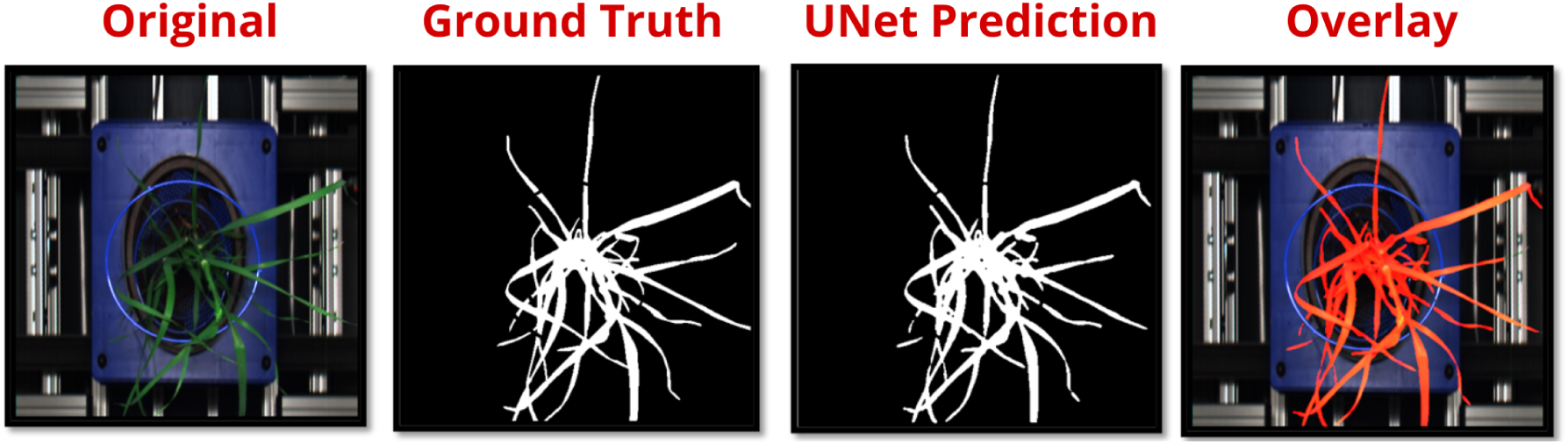
Comparison of U-Net segmentation performance against ground truth. The original image (left) is followed by the ground truth mask, the U-Net predicted segmentation, and an overlay of the prediction on the original image, highlighting segmentation accuracy and potential discrepancies.

To further validate the effectiveness of the BSA’s segmentation framework, we compared its performance against several established segmentation algorithms, including Otsu’s thresholding, K-means clustering, fuzzy C-means clustering, LinkNet, and DeepLabv3 [37–41]. The results, as shown in Figure 8, reveal notable differences in segmentation quality across each of these algorithms chosen for evaluation.

**Figure 8:**
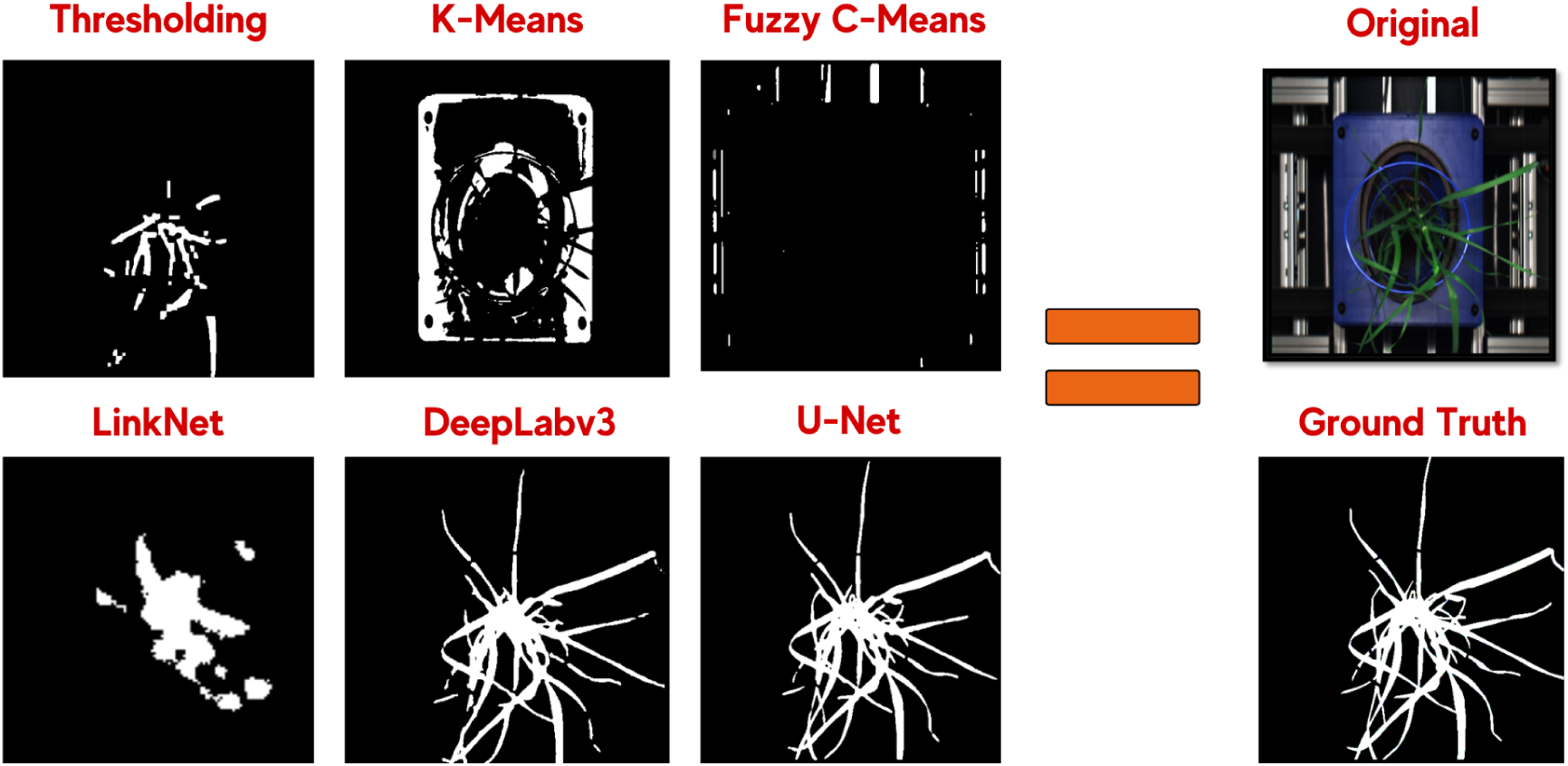
Comparison of segmentation methods for plant extraction. Traditional approaches (Thresholding, K-Means, Fuzzy C-Means) and deep learning-based models (LinkNet, DeepLabv3, U-Net) are evaluated against the ground truth. The original image is shown for reference, demonstrating the effectiveness of different segmentation techniques.

Thresholding-based algorithms, such as Otsu’s method, exhibited significant fragmentation and inconsistencies, highlighting the algorithms limitations in handling complex plant imagery. Current studies suggests that traditional thresholding-based segmentation algorithms in CV often struggle with accurately segmenting plant imagery, primarily due to their sensitivity to noise from background elements such as machinery and soil [42]. As a result, these algorithms frequently produce highly fragmented segmentations of RGB and hyperspectral images [43, 44]. Our findings corroborate these challenges, as Otsu’s method exhibited significant fragmentation and inconsistencies in boundary delineation.

In contrast, clustering-based algorithms, such as K-means and fuzzy C-means, performed slightly better but still resulted in significant detail loss. Clustering-based algorithms have shown previous promise in plant segmentation tasks [43]; however, their reliance on manual feature engineering and present cluster centroids can impede their ability to capture the full complexity of plant phenotypes [39, 45].

Among DL-based algorithms tested, LinkNet demonstrated a moderate segmentation performance but failed to achieve the fine-grained accuracy required for precise plant image segmentation. While LinkNet has previously demonstrated strong results in both remote sensing and medical imaging [35, 46], our findings indicate that its segmentation masks produced in this study were incomplete, likely due to its relatively shallow DL architecture. DeepLabv3, an advanced DL model designed for multi-scale segmentation [47], produced more coherent segmentation results than LinkNet. However, our evaluation of this algorithm revealed that despite the algorithms strengths, DeepLabv3 introduced segmentation inconsistencies into each result. These inconsistencies were particularly noticeable in hyperspectral images of fully grown plants, a challenge previously highlighted by Smith et al. (2021), who reported similar segmentation challenges with DeepLabv3 in dense plant canopies. When examining the accuracy of the BSA’s segmentation framework, we observed a marked improvement over the previously discussed algorithms. Unlike LinkNet, which exhibited difficulties with fine-grained segmentation, or DeepLabv3, which, despite its advanced multi-scale processing, introduced inconsistencies in hyperspectral imagery, the BSA consistently delivered precise and coherent segmentation results. Its ability to effectively segment plant images, even in challenging conditions such as overlapping foliage in hyperspectral imagery and variable lighting, underscores its robustness for image segmentation.

Although these results demonstrate the effectiveness of the BSA’s segmentation framework, further work is required to assess its generalisability across diverse imaging conditions. As highlighted in current studies, image segmentation in field-based environments is a major challenge given the variability introduced by non-standardised conditions [43, 44]. This includes challenges such as varying lighting conditions, occlusions, atmospheric interference, and differences in spatial resolution, making field-based segmentation more complex than in controlled environments [44]. Unlike the standardised conditions of a PSI imaging suite, data collected from ground-based sensors, UAVs, and satellites inherently introduce these variations, further complicating segmentation tasks [44]. Furthermore, DL algorithms trained solely on controlled datasets may require augmentation strategies or transfer learning techniques to effectively adapt the algorithm to combat these challenges [48].

Understanding these challenges requires a closer examination of the specific limitations associated with different imaging platforms. Ground-based sensors, including proximal phenotyping cameras and handheld spectral devices, provide high-resolution imaging but are often limited by inconsistent angles, shadows, and background noise from soil, water, or other non-plant elements [49]. Unmanned aerial vehicles (UAV-based imaging), on the other hand, offers flexible, high-throughput data collection over large agricultural fields but is sensitive to weather conditions, flight altitude, and wind-induced distortions [50, 51]. Finally, satellite imagery, while valuable for large-scale crop monitoring, typically suffers from lower spatial resolution, cloud cover interference, and temporal constraints due to revisit times [52].

Given these challenges, future work on the BSAs segmentation framework should explore domain adaptation techniques to enhance the framework’s applicability across different imaging environments. One potential approach involves using synthetic data augmentation strategies to introduce realistic variations in illumination, perspective, and occlusions [53]. Unlike other data augmentation techniques, synthetic data generation allows for precise control over these factors by creating tailored datasets that mimic real-world imaging conditions with high fidelity [54]. This level of control enables more diverse and representative training data, improving the framework’s robustness across different environments.

Furthermore, employing self-supervised learning and transfer learning methods could enhance the performance of the DNNs [55]. This could be achieved by fine-tuning the DNNs using smaller subsets of annotated field-based datasets, while leveraging large-scale remote sensing datasets to improve the generalizability of the segmentation framework. An approach demonstrated by Manas et al. (2021), who showed that pretraining on extensive remote sensing data (Sentinel-2) significantly boosts model adaptability to new imaging environments [56]. Furthermore, exploring multi-modal data fusion techniques that integrate spectral, thermal, and LiDAR-based (Light Detection and Ranging) segmentation may enhance accuracy across diverse sensing modalities, a common approach in self-driving car research [57].

In summary, our BSA segmentation framework demonstrates substantial improvements over both traditional and modern segmentation algorithms in controlled conditions. Its strong performance is due to its optimised U-Net design, which helps to capture spatial patterns while preserving important details needed for accurate segmentation. These findings support the growing consensus that DL-based segmentation models, particularly U-Net variants, offer significant advantages over traditional algorithms in plant phenotyping and other high-precision imaging applications, often making them the preferred choice among researchers.

### 4.2 BSA Excels in Multi-Species Plant Image Segmentation and Analysis

The selected case studies serve as a critical benchmark for assessing the effectiveness of our BSA segmentation framework in multi-species plant image segmentation and analysis using a set of imaging sensors. By applying the BSA segmentation framework to three distinct datasets — each incorporating different plant species, imaging sensors, and environmental conditions — we provide a comprehensive evaluation of its robustness, accuracy, and adaptability. This multi-dimensional validation is essential for demonstrating the broader applicability of the BSA in plant phenotyping and related fields, where image segmentation models must generalize across diverse biological and technical challenges.

The segmentation results depicted in Figure 9 demonstrate the results for the three species examined — wheat, barley, and Arabidopsis — demonstrating the BSA’s capacity to accurately segment plant imagery across diverse morphological characteristics and imaging conditions. The segmentation results generated by the BSA were highly consistent with the ground truth images across the three evaluated datasets, which further substantiates the effectiveness of the DNN incorporated into the BSA framework.

**Figure 9:**
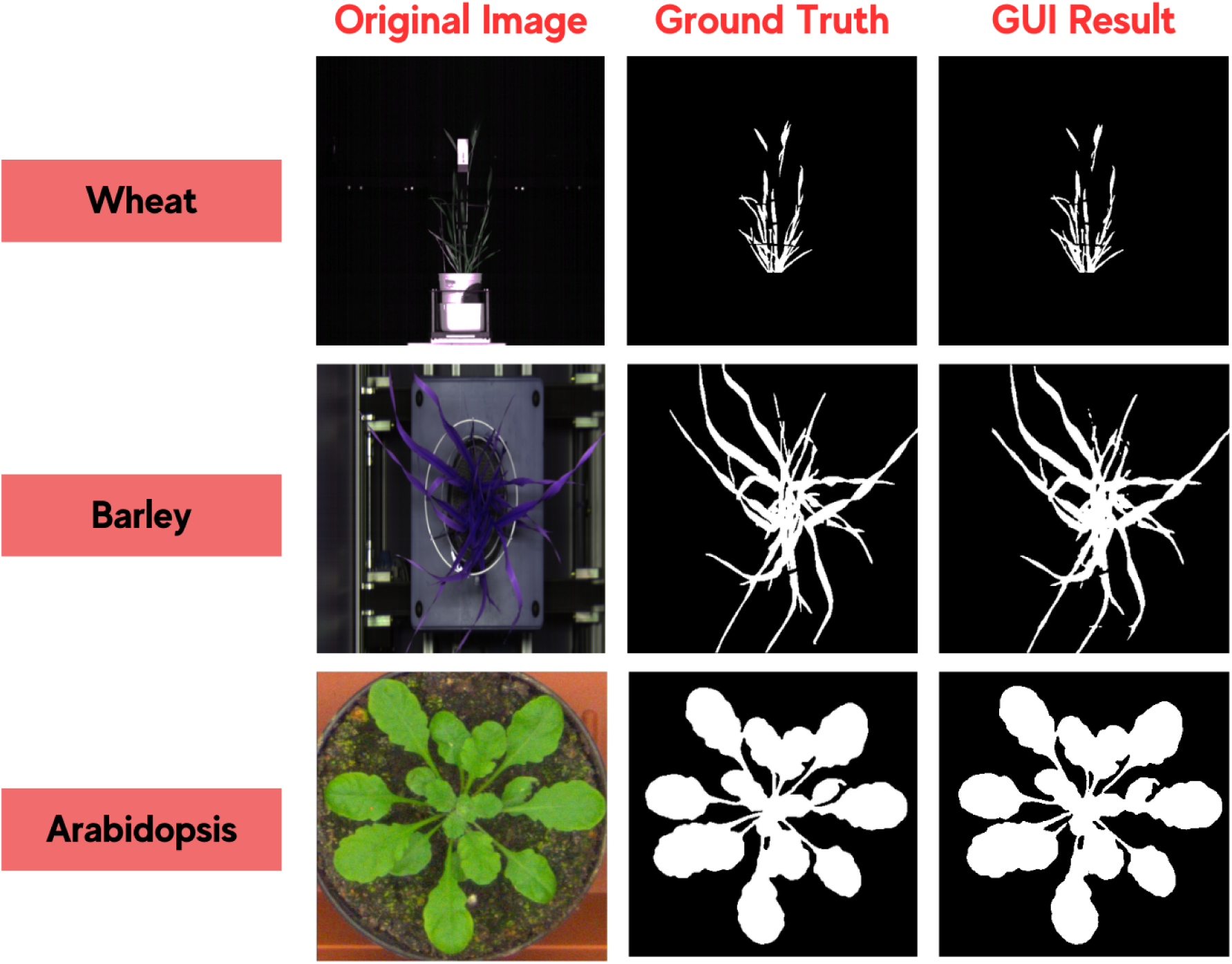
Segmentation results for different plant species using the Botanic Spectrum Analyser (BSA) GUI. The first column shows the original images of Wheat, Barley, and Arabidopsis. The second column presents the ground truth segmentation masks, while the third column displays the segmentation results generated by the BSA GUI, demonstrating accurate plant extraction across different species.

Across all three datasets, the BSA achieved an average accuracy of 99.7%, with an F1-score consistently exceeding 98% for each result, underscoring the precision of the final DNN integrated into the BSA. The Jaccard index and recall performance metrics further confirmed strong agreement with the ground truth images, which reinforces the robustness of the BSAs ability to capture plant features across multiple conditions.

For the wheat dataset, the BSA achieved a high segmentation accuracy (°99.5%), with F1 scores ranging between 81% and 89% across each of the sample predictions. While performance remained strong, slight variability in Jaccard index values suggested that factors such as illumination changes, occlusion, and overlapping leaf structures posed challenges for achieving a more precise segmentation result with this data [58].

These segmentation results are consistent with research that highlight the difficulty of segmenting dense cereal crops, where leaf curvature, leaf overlap and self-shading complicate image processing algorithms [59, 60]. Zhang et al. (2022) addressed these challenges by developing Wheat-Net, an instance segmentation model designed to detect wheat spikes in complex field conditions. Their model demonstrated robustness against occlusion, background noise, and variable illumination, achieving an average precision of 90% for mask segmentation and 99.29% accuracy in wheat spike counting. Nonetheless, high recall values suggest that the BSA segmentation framework successfully identifies the relevant plant features, even in the complex context of 2D hyperspectral imagery.

The segmentation results for the barley dataset closely mirrored those of wheat, demonstrating the BSA’s ability to generalize across different cereal crops. Despite differences in leaf architecture and growth patterns, the framework maintained high accuracy and F1 scores, reinforcing its effectiveness in handling crop-to-crop variability. This consistency is particularly noteworthy given that traditional rule-based segmentation algorithms often struggle with species-specific variations in texture and reflectance properties [44]. For example, Arroyo et al. (2016) found that thresholding-based methods, such as Otsu’s approach, were highly sensitive to illumination changes and variations in plant growth stages, leading to inconsistent segmentation results. Their study demonstrated that machine learning-based alternatives, such as k-Nearest Neighbors (k-NN), were more robust under diverse field conditions.

Finally, in Arabidopsis, a model plant species, the BSA segmentation framework was able to effectively segment each individual plant, despite its smaller size and more intricate structural features. Further observations revealed that the model consistently achieved an accuracy exceeding 99% for each segmentation, underscoring its capability to discern even the finest leaf edges from the surrounding background with enhanced sensitivity. Despite this high accuracy, Arabidopsis plants present a relatively straightforward segmentation task compared to other crop species. Its 2D architecture, small size and well-defined leaf structures contribute to higher segmentation accuracy, particularly in controlled environments where occlusion and background complexity are minimal [61]. Traditional segmentation techniques, such as thresholding-based methods, have successfully achieved high accuracy in Arabidopsis segmentation [62, 63]. For example, Henke et al. (2021) employed a k-means clustering method (kmSeg) that achieved segmentation accuracy between 96–99% for Arabidopsis, highlighting the effectiveness of non-deep learning approaches in structured image environments [64]. Similarly, Hüther et al. (2020) demonstrated that colour-based thresholding methods could reliably segment Arabidopsis rosettes in phenotyping experiments, though performance declined when plants exhibited non-standard colouration or background variations [65]. These findings suggest that while deep learning approaches enhance segmentation under complex conditions, classical methods can still be effective for specific plant species such as Arabidopsis.

These findings confirm the versatility and robustness of the BSA segmentation framework across diverse plant species and imaging conditions. The framework consistently delivered high segmentation accuracy for wheat, barley, and Arabidopsis, demonstrating its adaptability to varying morphological structures and technical constraints. Its ability to generalize across different datasets (which included RGB, VNIR and SWIR data) underscores its potential for automated plant phenotyping, precision agriculture, and crop monitoring. By effectively addressing the challenges posed by species-specific variations and environmental factors, the BSA segmentation framework represents a reliable tool for plant image analysis in both research and applied agricultural settings.

### 4.3 BSA’s Advanced Capabilities in Root System Analysis

We also tested the BSA’s adaptability for segmenting plant root images. Roots are inherently complex structures, notoriously challenging to segment accurately due to their intricate morphology and overlapping patterns [66]. Like plant images, root imagery typically exhibits lower contrast and higher background noise, making segmentation tasks significantly more difficult. Moreover, inadequate segmentation of root architecture has substantial downstream effects on root phenotyping. Inaccurate segmentation can lead to distorted measurements of critical traits such as total root length, branching patterns, and growth angles, compromising the reliability of phenotypic analyses [66].

Despite this, several solutions have been developed for root segmentation in conventional RGB images, leveraging well-established image processing techniques and ML frameworks [67–69]. However, the segmentation of roots using hyperspectral imaging remains an area with limited research [66]. Unlike RGB imaging, hyperspectral data offers richer spectral information. However, the complexity of analysing such high-dimensional datasets presents unique computational and analytical challenges. Consequently, robust methods tailored explicitly for hyperspectral root image segmentation are still scarce, highlighting an important gap and opportunity for future research. To further explore the effectiveness of the BSA, we assessed its capability for root image segmentation without any additional retraining specific to root images. The rationale behind avoiding retraining the BSA was to evaluate the network’s existing applicability and robustness for this task before any future work could be assigned to it. Despite the absence of domain-specific retraining, the results from the BSA demonstrated remarkable accuracy in root image segmentation.

Additional experiments were conducted to further support these findings and benchmark the BSA against current commercial segmentation algorithms. As illustrated in Figure 10, the segmentation results achieved by the BSA were distinctly superior. Current commercial segmentation algorithms exhibited suboptimal, suggesting promising avenues for future optimisation. Quantitative performance analysis (shown in Table 2) further reinforced these observations, indicating a 50% improvement in the F1-score for the BSA compared to the commercial segmentation solution tested. The commercial segmentation algorithms evaluated in below-ground segmentation demonstrated certain limitations in their ability to robustly handle the intricate structural complexity characteristic of root systems. The overlapping structures characterising root architectures, including fine lateral branches and inconsistent contrast against diverse backgrounds, pose substantial challenges for traditional segmentation algorithms [66]. Consistent with previous literature, this underperformance highlights that classical, traditional segmentation algorithms (e.g., thresholding and edge detection) tend to be less effective when applied to complex images such as roots [70]. Even when using ML-based commercial tools, these models typically focus on optimising contrast for well-separated objects and struggle to generalise to scenarios involving fine-grained textures and low signal-to-noise ratios, such as root systems situated within soil or other complex media [68, 69].

**Figure 10:**
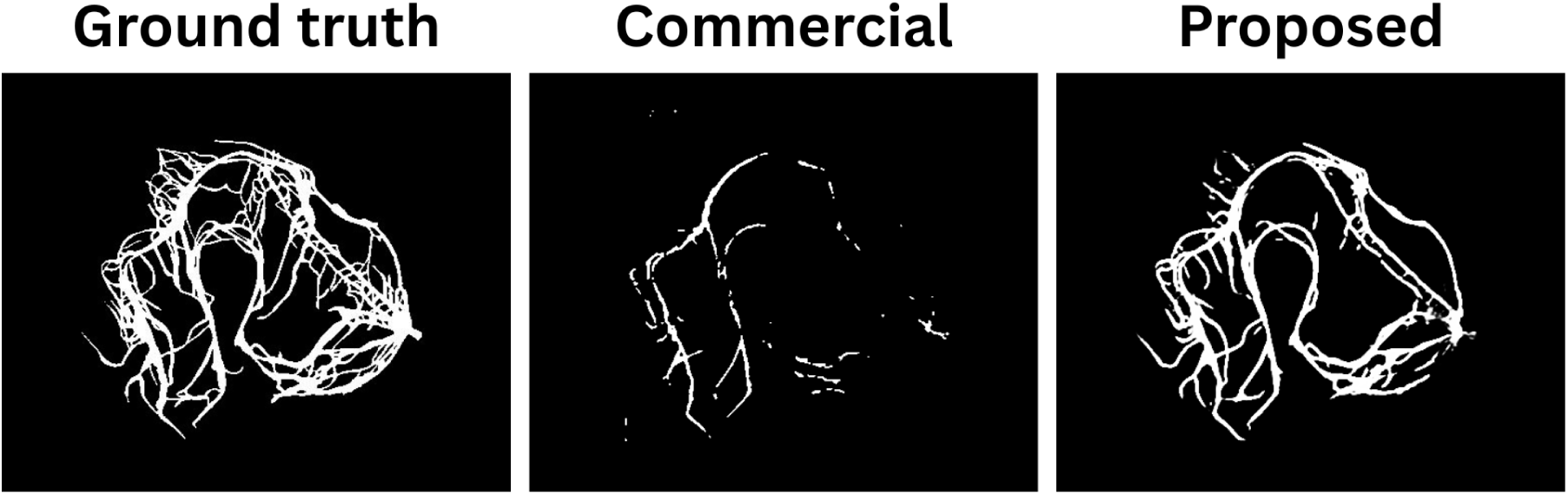
Segmentation results of hyperspectral root images. From left to right: Ground truth annotation, output from a commercial segmentation tool, and result from the Botanical Spectrum Analyser (BSA). The BSA demonstrates superior structural continuity and accuracy in capturing fine root details compared to the commercial approach.

**Table 2:**
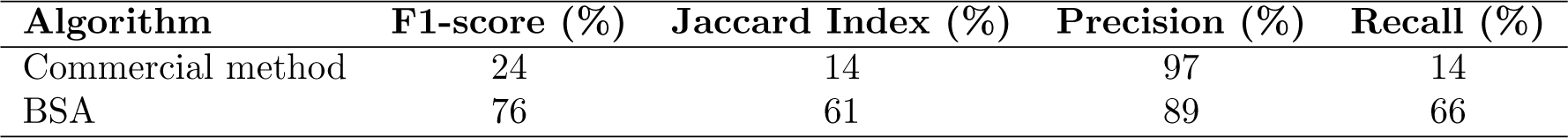
Quantitative comparison of segmentation performance between the commercial method and the proposed Botanical Spectrum Analyser. The proposed method achieves significantly higher F1-score, Jaccard Index, and Recall, while maintaining high Precision, indicating more balanced and accurate root segmentation.

An important consideration from our BSA GUI evaluation for root image segmentation is the influence of the imaging modality. In our experiments, root images were acquired using a top-down angle with a SWIR imaging sensor, which provided consistent illumination and uniform backgrounds. This favourable imaging setup significantly enhanced the BSA’s segmentation capability, contributing to its notable accuracy despite the absence of domain-specific retraining.

However, exploratory trials using root images acquired in soil conditions yielded significantly poorer results, indicating a pronounced dependency of segmentation accuracy on imaging background. Soil introduces additional complexities such as heterogeneous textures, varying moisture levels, and reduced contrast between roots and the surrounding medium, substantially increasing segmentation difficulty [66]. Literature confirms that such environmental variations significantly influence segmentation performance, particularly when fine structures and complex overlapping are involved [71]. These in-soil images were captured using the same PlantScreen modular system, SWIR camera, and imaging angle as the controlled trials, underscoring the importance of developing models robust to real-world imaging conditions.

Overall, the evaluation of the BSA for root image segmentation highlights its promising capabilities in addressing the unique challenges posed by complex root structures, even without domain-specific retraining. Despite some remaining areas for improvement, the substantial performance advantage over existing commercial tools underscores the need for more specialised approaches in root phenotyping. Future work should focus on fine-tuning models like the BSA and expanding annotated root image datasets to unlock even greater segmentation accuracy. Advancements in this area have the potential to significantly enhance phenotypic analyses, with broad implications for agricultural innovation, ecological research, and plant science.

### 4.4 BSA Stands Out with User-Friendly Design for Non-Technical Users

One of the primary challenges in computational plant phenotyping is accessibility [72, 73]. Many non-expert users, including plant biologists, agronomists, and ecologists, face significant technical barriers when working with advanced image analysis software. Traditional tools often require proficiency in CV, computer programming, or data analytics, which limits their usability for researchers without formal computational training. Recognizing this limitation, the BSA GUI was developed to democratize advanced image analysis, ensuring that researchers can leverage DL-based segmentation without requiring extensive technical expertise.

However, existing image analysis tools, while powerful, often reinforce these accessibility challenges. For example, OpenCV, one of the most well-established computer vision libraries, offers a comprehensive suite of tools for image segmentation, feature extraction, and object recognition [74]. Due to its efficiency and versatility, OpenCV has been widely adopted across various scientific fields, including plant phenotyping for a variety of image analysis tasks [75–77]. However, OpenCV is primarily designed for computer vision specialists who are proficient in programming and algorithm development. Despite its invaluable capabilities, its reliance on scripting and coding presents a barrier for researchers without experience in Python or C++.

To address the need for plant-specific image analysis tools, PlantCV was developed as an extension of OpenCV, tailored specifically for plant science applications. It provides functions for trait quantification, colour segmentation, and feature extraction, making it a powerful tool for high-throughput phenotyping studies. PlantCV has been widely used to quantify leaf morphology, detect stress symptoms, and analyse growth patterns [78–80]. However, despite its domain-specific focus, PlantCV still requires users to write Python scripts or work within Jupyter Notebooks, limiting accessibility for researchers unfamiliar with coding. Although scripting-based workflows enhance reproducibility and flexibility, they pose usability challenges for biologists and agronomists without formal programming expertise. Research on the adoption of computational tools in biological sciences has highlighted that the lack of user-friendly interfaces remains a major barrier to integrating advanced data analytics into standard research workflows [81].

GUIs have been widely recognized as a key factor in improving the accessibility of computational tools for non-technical users [82]. A well-designed GUI can allow users to interact with powerful computational tools without requiring prior programming knowledge, thus lowering the entry barrier for image analysis in plant science. SeedExtractor, for instance, is an open-source GUI developed for seed image analysis [83]. The GUI enables users to efficiently extract seed traits such as size, shape, and colour through an intuitive interface, supporting multiple colour spaces and batch processing for high-throughput analysis. Its effectiveness has been validated across diverse plant species and in genome-wide association studies, demonstrating its reliability for phenotypic data extraction in plant science. However, SeedExtractor is highly specialized for seed trait analysis and lacks the flexibility to accommodate a broader range of plant phenotyping tasks, limiting its applicability to researchers working with different plant structures or complex segmentation problems.

Several other GUI-based tools have been developed to address specific needs in plant phenotyping. PhenoImage provides a user-friendly environment for analysing plant images with minimal coding requirements, integrating machine learning-based segmentation to facilitate trait extraction in high-throughput studies [84]. JustDeepIt offers a GUI-driven solution for deep learning-based image segmentation, allowing researchers to apply pre-trained deep learning models to plant images without requiring expertise in neural networks [85]. Although powerful, JustDeepIt focusses primarily on inference rather than model training, limiting customisation for specialised applications. Finally, RhizoVision, designed specifically for root phenotyping, provides an intuitive interface to analyse root architecture traits, making it particularly useful for studies on plant growth and soil interactions [67]. However, like SeedExtractor, RhizoVision is specialised for a narrow application domain, reducing its flexibility for broader plant phenotyping tasks.

In contrast, the BSA GUI was developed with the goal of making advanced image segmentation and hyperspectral image analysis accessible to all users, regardless of their technical background. Unlike OpenCV and PlantCV, which require coding proficiency, BSA offers a fully interactive GUI, allowing plant scientists to upload images, apply DL-based segmentation models, and analyse results without writing a single line of code. The GUI provides the following key features:

1. Intuitive design: The BSA interface follows best-practices in GUI design to ensure a seamless user experience [86]. Features such as drag-and-drop image loading, real-time visualisation of results, and adjustable segmentation parameters enhance interactivity and user engagement.
2. Automated DL integration: The framework incorporates pre-trained DNNs optimised for plant segmentation which eliminates the need for users to train their own models from scratch. This is consistent with recent developments in automated machine learning tools (e.g., AutoML [87]), which seek to democratize AI by lowering the expertise required to leverage complex algorithms.
3. Multi-sensor support: Unlike existing tools that primarily focus on either RGB or hyperspectral imaging, BSA supports multiple imaging sensors, including RGB, VNIR, and SWIR, enabling researchers to analyse a broader range of plant traits.
4. Cross-disciplinary usability: By eliminating coding requirements, BSA fosters interdisciplinary collaboration, enabling plant scientists, agronomists, and ecologists to conduct high-quality image analysis independently.

The development of GUI-based tools has significantly improved the accessibility of computational plant phenotyping, addressing long-standing technical barriers faced by researchers without any programming expertise. While existing tools like OpenCV, PlantCV, and specialised GUIs such as SeedExtractor and RhizoVision offer valuable functionalities, they often remain limited by either their technical complexity or narrow application scope. The BSA GUI was designed to overcome these challenges by providing an intuitive, fully interactive platform for advanced image segmentation and hyperspectral image analysis. By integrating DL algorithms and supporting multiple imaging sensor types, the BSA democratizes plant phenotyping, enabling researchers across disciplines to conduct high-quality image analysis without requiring extensive computational training. This approach represents a step toward broader adoption of AI-driven image analysis in plant science, fostering interdisciplinary collaboration and expanding the potential for data-driven discoveries in agricultural and ecological research.

## 5 Conclusion

In this study, we introduced the BSA, a user-friendly graphical interface designed to simplify and enhance hyperspectral and RGB image segmentation for plant phenotyping. Using a deep neural network (DNN)-based framework, the BSA automates segmentation with high accuracy, minimising the need for technical expertise. Across diverse case studies involving wheat, barley, and Arabidopsis using RGB, VNIR, and SWIR shoot imaging, the framework consistently achieved F1 scores exceeding 98%, outperforming traditional and DL-based methods.

Beyond accuracy, the BSA’s intuitive, tab-based interface and structured workflow improve accessibility for non-experts, enabling seamless integration with various sensors and species. This broad applicability spans controlled environments to large-scale agricultural monitoring. Key features include automated segmentation, multi-sensor support, and usability for researchers without computer vision backgrounds.

We also evaluated the BSA for root image segmentation, which is typically a complex task due to low contrast and structural complexity. Without retraining, the BSA outperformed commercial tools in top-down SWIR images, improving F1 scores by up to 50%. However, the images of roots in the soil led to less consistent results, indicating the need for a targeted adaptation of the model. Future work should explore the BSA’s scalability in field settings, where variability and image noise present further challenges. Enhancements like domain adaptation, advanced transfer learning, and multi-modal fusion could improve robustness. Cloud or mobile deployment could also support the broader adoption of high-throughput and precision agriculture.

Overall, the BSA bridges the gap between advanced deep learning and practical plant science, offering a scalable solution for automated image analysis. Its continued development will support the growing need for accessible high-throughput phenotyping tools in modern agriculture.

## Acknowledgements

None

## Author Contributions

Conceptualization: JJW and LG; data curation and visualisation: JJW, EJ, VP, and LG; writing original draft: all authors; writing, reviewing, and editing: JJW and SN; funding acquisition: SN.

## Funding

This research was supported by Science Foundation Ireland Centre by the SFI President of Ireland Future Research Leaders to SN under Grant No. 18/FRL/6197.

## Conflicts of Interest

The authors declare no conflict of interest.

## Data Availability

The botanical spectrum analyser graphical user interface and a sample of the data used for testing purposes are available for download from the following webpage: https://csi-dublin.ie/

## Supplementary Materials

Not applicable.

## Notes

https://csi-dublin.ie

### Competing Interest Statement

The authors have declared no competing interest.

